# Metabolic reprogramming of hypoxic tumor-associated macrophages through CSF-1R targeting favours treatment efficiency in colorectal cancers

**DOI:** 10.1101/2024.02.22.581558

**Authors:** Khaldoun Gharzeddine, Cristina Gonzalez Pietro, Marie Malier, Clara Hennot, Renata Grespan, Yoshiki Yamaryo-Botté, Cyrille Y. Botté, Fabienne Thomas, Marie-Hélène Laverriere, Edouard Girard, Gael Roth, Arnaud Millet

## Abstract

Tumor-associated macrophages participate in the complex network of support that favors tumor growth. Among the various strategies that have been developed to target these cells, blockade of the CSF-1R receptor is one of the most promising one. Here, we characterize the resulting state of human macrophages exposed to a CSF-1R kinase inhibitor. We find that CSF-1R receptor inhibition in human macrophages is able to impair cholesterol synthesis, fatty acid metabolism and hypoxia-driven expression of dihydropyrimidine dehydrogenase, an enzyme responsible for the 5-fluorouracil macrophage-mediated chemoresistance. We show that this inhibition of the CSF-1R receptor leads to a downregulation of the expression of SREBP2, a transcription factor that controls cholesterol and fatty acid synthesis. We also show that the inhibition of ERK1/2 phosphorylation resulting from targeting the CSF-1R receptor destabilizes the expression of HIF2α in hypoxia resulting in the downregulation of dihydropyrimidine dehydrogenase expression restoring the sensitivity to 5-fluorouracil in colorectal cancer. These results reveal the unexpected metabolic rewiring resulting from the CSF-1R receptor targeting of human macrophages in tumors.

## Introduction

A large body of work has been reported providing compelling evidence that tumor-associated macrophages (TAMs) are involved in tumor growth, treatment resistance and metastasis (DeNardo & Ruffell, 2019; Engblom *et al*, 2016). Indeed, TAMs are associated with poor prognosis in the vast majority of solid tumors (Cortese *et al*, 2020). Consequently, great efforts have been made to target TAMs in order to direct the immune response against the tumor (Mantovani *et al*, 2022). CSF-1R is a receptor with intrinsic tyrosine kinase activity that recognizes CSF-1 (M-CSF, macrophage colony stimulating factor) and IL-34. The CSF-1/CSF-1R signaling pathway is involved in macrophage survival, proliferation, differentiation and chemotaxis (Stanley & Chitu, 2014). This pivotal role opens the way to use CSF-1R receptor targeting in various pathological contexts and especially in cancer to modulate macrophages (Cannarile *et al*, 2017). CSF-1R receptor targeting has been used in various tumor mouse models with promising results but only modest beneficial effects in humans when used alone (Mantovani *et al*, 2022). Furthermore, the recognition that the type of tumor model used (O’Brien *et al*, 2021) and the modification of the cellular environment such as oxygen availability (Pradel *et al*, 2016), affect the efficiency of the CSF-1R targeting highlights the need for a better understanding of the functional consequences of CSF-1R receptor targeting on human macrophages.

The importance of CSF-1R receptor blockade beyond the induction of apoptosis has been recognized in animal models that advocate the possibility of achieving an anti-tumor phenotypic reprogramming (Pyonteck *et al*, 2013). However, the mechanisms involved in macrophage’s reprogramming by CSF-1R targeting are unclear.

Here, we characterize the functional consequences of CSF-1R receptor inhibitors on human macrophages. We show that CSF-1R receptor inhibition is associated with impaired cholesterol and fatty acid synthesis under the control of the transcription factor SREBP2. Furthermore, CSF-1R receptor blockade leads to an inhibition of the phosphorylated state of ERK1/2 resulting in a decrease in the expression of HIF2α, a hypoxic response transcription factor. A consequence of the HIF2α-dependent hypoxic response is the downregulation of dihydropyrimidine dehydrogenase expression, the limiting rate enzyme of the first step of the pyrimidine catabolic pathway, which is responsible for a hypoxia-driven macrophage-mediated chemoresistance to 5-fluorouracil in colorectal cancer (Malier *et al*, 2021).

## Results

### Complete CSF-1R inhibition is associated with a profound phenotypic change without death induction in human macrophages

Upon ligation of CSF-1, the CSF-1 receptor dimerizes, resulting in multiple site phosphorylation of the intracellular tail of the receptor (Stanley & Chitu, 2014). To investigate the resulting effect of inhibition of the CSF-1R receptor-dependent signaling pathway, we first evaluated the response of tyrosine 723 phosphorylation to increasing doses of edicotinib, a selective CSF-1R tyrosine kinase inhibitor (Kumar *et al*, 2017; von Tresckow *et al*, 2015; Genovese *et al*, 2015). We found that edicotinib at 10µM is able to completely inhibit CSF-1R’s phosphorylation (Figure 1A). Targeting the CSF-1R receptor with tyrosine kinase inhibitors is a strategy used in mice to deplete macrophages (Ries *et al*, 2014). However, we found that the complete inhibition of CSF-1R phosphorylation in human primary macrophages was not associated with cell death induction (7AAD and Annexin V) (Figure 1B & Supplementary Figure 1) or disruption of cellular metabolic activity (MTT) (Figure 1C). The importance of CSF-1 in macrophage differentiation and polarization prompted us to evaluate the effect of CSF-1R receptor blockade on human macrophage phenotype. To achieve this goal, we performed a global gene expression analysis by RNA sequencing (RNAseq) on bulk macrophages 48h after exposure to vehicle and edicotinib. To extract differentially expressed genes between treated and naïve macrophages, we used a moderated t-test with an adjusted p-value (Benjamini-Hochberg procedure with a threshold set at q-value <0.05) and a fold change threshold set at 2 (log_2_ FC <-1 or Log_2_ FC > 1). CSF-1R receptor inhibition by edicotinib was associated with 740 downregulated and 481 upregulated genes (Figure 1D). We then used a Gene Ontology (GO) enrichment analysis to identify pathways affected by edicotinib. The GO terms associated with genes downregulated by edicotinib indicated that the regulation of the cholesterol metabolic pathway, the inhibition of the ERK1/2 pathway, and lipid synthesis were the main of biological processes identified as a result of CSF-1R receptor blockade (Figure 1E).

**Figure 1.**
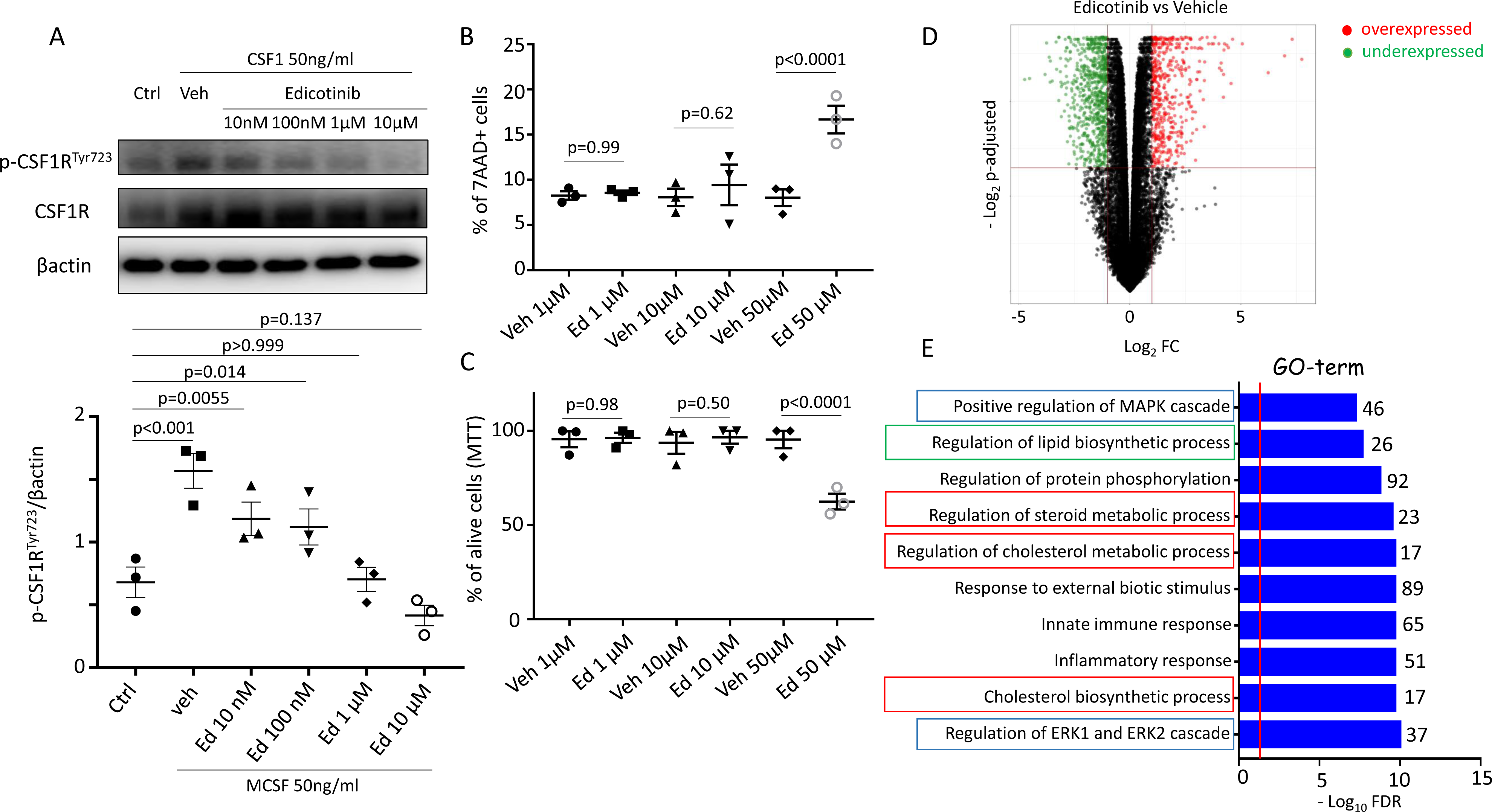
Complete CSF-1R inhibition is associated with a profound phenotypic change. (A) Immunoblot of phosphorylated CSF-1R at site Tyrosine 723, CSF-1R total protein and β-actin (upper panel). Quantification of the level of phosphorylation normalised to β-actin from three independent experiments. (B) Flow cytometry analysis of 7-AAD+ cells treated for 48h. Each condition has its own vehicle control (DMSO) at the same concentration than edicotinib (Ed) (n=3). (C) MTT assay on cells exposed to Edicotinib for 48h (n=3). (D) Volcano-plot analysis of differentially expressed genes between edicotinib vs vehicle (n=4). (E) GO (Gene Ontology)-term analysis of biological processes associated with downregulated genes under edicotinib exposure. Number of genes associated with each GO-terms are indicated.

### CSF-1/CSF-1R regulates cholesterol synthesis in human macrophages via the transcription factor SREBP2

To explore the gene ontology identification of cholesterol synthesis regulation under CSF-1R targeting, we examined the expression of all the enzymes involved in the cholesterol synthesis starting from acetyl-CoA. Cholesterol, an essential lipid component of mammalian cells, is mainly obtained by endogenous synthesis or uptake. Endogenous synthesis relies on nearly 30 enzymatic reactions (Nes, 2011). We found that genes coding for enzymes involved in cholesterol synthesis *ACAT2*, *HMGCS1, HMGCR, MVK, PMVK, MVD, IDI1, FDPS, FDFT1, SQLE, LSS, CYP15A1, TMF7SF2, NSDHL, HSD17B7, MSMO1, SC5D, EBP, DCHR7, DCHR24* are downregulated under edicotonib exposure (Figure 2A) and except for two of them (*PMVK*, *LSS*) the fold change is greater than 2 (Figure 2B). The control of cholesterol synthesis genes by the CSF-1/CSF-1R pathway was confirmed by the stimulation of human macrophages by CSF-1, which showed the upregulation of *HMGCR* and *DHCR7* (Figure 2C). SREBP2 (Sterol regulatory element-binding protein 2) is a helix-loop-helix leucine zipper transcription factor that regulates the synthesis and cellular uptake of cholesterol (Horton *et al*, 1998). Using the ENCODE data set (The ENCODE (ENCyclopedia Of DNA Elements) Project, 2004), we have explored the known targets of this transcription factor and found that the cholesterol synthesis genes are under the control of SREBP2 (Figure 2D). Interestingly, the LDL receptor (*LDLR*) which is responsible for cellular uptake of cholesterol from the extracellular milieu and the fatty acid synthetase (*FASN*) are also SREBP2 target genes (Figure 2D). We validated the SREBP2 dependent expression of cholesterol synthesis genes (*HMGCR*, *DHRC7*) and *LDLR* using RNAi targeting against SREBF2, the gene encoding the SREBP2 protein (Figure 2E). We demonstrated the control of *SREBF2* by the CSF-1/CSF-1R pathway by showing that *SREBF2* expression is increased under CSF-1 stimulation (Figure 2F) and decreased when CSF-1R is blocked by edicotinib (Figure 2G). We confirmed that the active form of SREBP2 expression is downregulated by the CSF-1R blockade (Figure 2H). In addition, *LDLR*, a target of SREBP2, is downregulated by edicotinib (Figure 2I). We observed the resulting decrease in cellular cholesterol levels under edicotinib exposure similar to what is found using RNAi targeting of SREBF2 or blockade of HMGCR activity with simvastatin, a specific inhibitor (Figure 2J). The cellular homeostasis of cholesterol is a balance between synthesis, uptake and efflux. The impairment of cholesterol synthesis and uptake by CSF-1R receptor blockade could be compensated by a decrease in cholesterol efflux. To evaluate this possibility, we studied the expression of the two major cholesterol efflux transporters expressed in macrophages ABCA1 and ABCG1. We found that the CSF-1R targeting did not decrease the expression of these transporters in macrophages (Supplementary Figure 2A) and that the decrease in intracellular cholesterol levels induced by CSF-1R receptor blockade resulted in a global decrease in macrophage cholesterol secretion (Supplementary Figure 2B).

**Figure 2.**
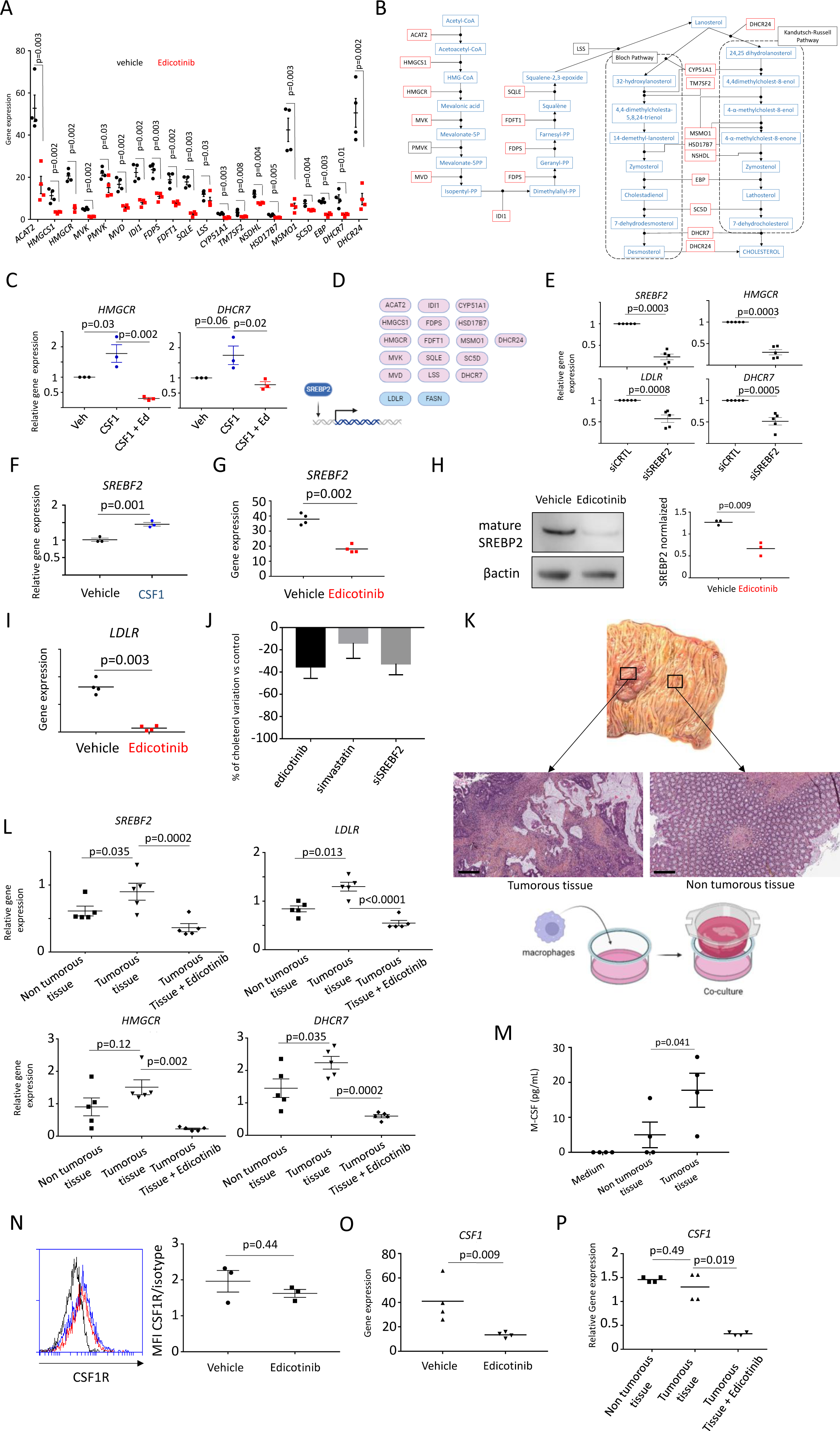
CSF-1/CSF-1R controls cholesterol synthesis in human tumor-associated macrophages. (A) Gene expression of cholesterol synthesis enzymes in human macrophages treated by edicotinib (in red) compared to controls (vehicle in black) (n=4). Gene expression levels are expressed in FPKM=Fragments Per Kilobase Million. (B) The cholesterol synthesis pathway with intermediate products. Enzymes which present a downregulation with a fold change >2 are represented in red. (C) Expression of *HMGCR* and *DHRC7* genes after 6h stimulation by CSF-1 50 ng/mL on previously starved human macrophages determinded by RT-qCPR (n=3). (D) Representation of SREBP2 target genes based on ChIP analysis (ENCODE study). (E) Expression of *SREBF2*, *HMGCR*, *DHCR7* and *LDLR* genes in RNAi treated macrophages against *SREBF2* determined by RT-qPCR (n=5). (F) Expression of *SREBF2* gene after 6h stimulation by CSF-1 50 ng/mL on previously starved human macrophages determinded by RT-qCPR (n=3). (G) Gene expression of *SREBF2* in human macrophages treated by edicotinib (n=4). Gene expression levels are expressed in FPKM=Fragments Per Kilobase Million. (H) Representative immunoblot of mature SREBP2 in human macrophages treated with edicotinib compared to controls (Left panel) using actin as a control loading. Quantification of SREBP2 normalized to the control loading is represented in the right panel (n=3). (I) Gene expression of *LDLR* in human macrophages treated by edicotinib (n=4). Gene expression levels are expressed in FPKM=Fragments Per Kilobase Million. (J) Quantification of intracellular cholesterol in macrophages treated by edicotinib or simvastatin and in RNAi against SREBF2 treated macrophages compared to vehicle (DMSO) or RNAi control respectively (n=3). The amount of decrease is represented in % of the appropriate control (DMSO for Edicotinib and simvastatin; siCTRL for SiSREBF2). (K) Macroscopic view of a colon adenocarcinoma and HES staining of tumorous and non-tumorous tissues used in a co-culture with human macrophages. (L) Expression of *SREBF2*, *HMGCR*, *DHCR7* and *LDLR* genes in hypoxic macrophages cultured with non tumorous and tumorous tissues determined by RT-qPCR (n=5). Tumor associated macrophages were also treated by edicotinib. Relative expression compared to control macrophages in normoxia. (M) ELISA quantification of IL-34 and CSF-1 secreted by tumorous and non-tumorous tissues (n=4). (N) Flow cytometry quantification of membrane CSF-1R expression on macrophages treated by edicotinib compared to controls (DMSO) (n=3). (O) Gene expression of *CSF-1* in human macrophages treated by edicotinib (n=4). Gene expression levels are expressed in FPKM=Fragments Per Kilobase Million. (P) Expression of *CSF-1* gene in hypoxic macrophages cultured with non tumorous and tumorous tissues determined by RT-qPCR (n=4). Tumor associated macrophages were also treated by edicotinib. Relative expression compared to control macrophages in normoxia.

### Edicotinib interferes with the macrophage-tumor communication resulting in modulation of cholesterol synthesis in colorectal cancers

Membrane cholesterol efflux from macrophages has been reported to be involved in tumor growth (Goossens *et al*, 2019). This efflux relies on the ability of macrophages to express efflux transporters but also on the maintenance of their intracellular cholesterol pool based on synthesis and uptake. To further understand the importance of cholesterol synthesis in the cross-talk between macrophages and tumors, we performed a co-culture between human macrophages and tumor or non-tumor tissues from the same colon obtained from surgical resection of colorectal adenocarcinomas (Figure 2K). We observed that macrophages under the influence of tumor tissue upregulate their cholesterol synthesis genes as well as the cholesterol uptake receptor LDLR under the control of SREBF2 which is upregulated by the tumor (Figure 2L). We also demonstrated that edicotinib is able to counteract this tumor-driven metabolic reprogramming by interfering with cholesterol synthesis and uptake (Figure 2L). This communication between tumors and macrophages suggested the involvement of the CSF-1/CSF-1R pathway. We confirmed that tumors secrete significantly more CSF-1 than non-tumor tissues, which drives cholesterol synthesis and uptake in macrophages (Figure 2M). Nevertheless, we sought to verify whether edicotinib could also affect the expression of the CSF-1R receptor on the surface of macrophages but found no decrease in receptor following CSF-1R blockade (Figure 2N). Interestingly, we found that the ability of macrophages to produce their own CSF-1 is impaired in human macrophages exposed to edicotinib (Figure 2O). Even when tumors do not increase the autocrine-associated production in macrophages, edicotinib is able to downregulate this production in macrophages, enhancing the blockade of the CSF-1/CSF-1R pathway (Figure 2P).

### CSF-1R receptor targeting downregulates HIF2α expression in hypoxia through inhibition of ERK1/2 phosphorylation

Previous studies have shown that several environmental factors can modulate the cellular response to CSF-1R receptor targeting (Pradel *et al*, 2016). Among these factors, hypoxia is of particular interest because the tumor environment is associated with low oxygen availability (Wilson & Hay, 2011). To assess the interplay between hypoxia and CSF-1R receptor targeting, we performed the study of differentially expressed genes in hypoxia compared to normoxia and analyzed the impact of edicotinib on the hypoxic response (Figure 3A). The cell survival profile of CSF-1R receptor blockade in hypoxia was similar to what we found in normoxia (Supplementary Figure 3A). Heat-map analysis revealed that specific hypoxia-sensitive genes exhibited two distinct patterns: specific hypoxia-sensitive genes that maintained their expression profile under CSF-1R receptor blockade (clusters 1 & 3) and genes with a modified profile under CSF-1R receptor blockade (clusters 2, 4 & 5). Among the hypoxia upregulated genes (cluster 3), we found several genes *LDHA*, *NDRG1*, *P4HA1* and *SLC2A1* (Figure 3B), which are part of the HIF1α target gene set identified by ChIP enrichment analysis (Lachmann *et al*, 2010) suggesting that the HIF1α-dependent hypoxic response is mainly independent of the CSF-1R pathway. Of particular interest, we observed that the downregulation of some hypoxic genes is abolished by CSF-1R receptor blockade (cluster 2) with an increased expression profile in normoxia (Figure 3A). One of this gene is *EPAS1* (Figure 3C). The *EPAS1* gene encodes the transcription factor HIF-2α, which plays an essential role in the cellular response to low oxygen environments (McGettrick & O’Neill, 2020). The downregulation of *EPAS1* mRNA in hypoxia contrasts with the stabilization of HIF-2α protein in the same condition (Figure 3D). To understand the role of CSF-1R in the HIF-2α-dependent hypoxic response, we sought to elucidate the function of HIF-2α by studying *FLT1* (VEGFR1), a HIF-2α target gene (Sasagawa *et al*, 2018). Indeed, *FLT1* expression is increased in hypoxic macrophages (Figure 3E) under the control of HIF-2α transcriptional activity, as demonstrated by RNAi targeting of EPAS1 in hypoxic macrophages (Supplementary Figure 3B). The decrease of FLT1 (Figure 3E) and the corresponding increase of EPAS1 mRNA level upon CSF-1R receptor blockade by edicotinib in hypoxic macrophages (Figure 3C) supports a decrease of HIF-2α expression in hypoxia under CSF-1R receptor blockade, a pattern that we confirmed by immunoblot (Figure 3F). Interestingly, the activation of HIF-2α leading to its translocation to the nucleus and fixation on Hypoxic Response Elements (HRE) gene promotors is ERK1/2 dependent (Conrad *et al*, 1999). The identification of the ERK1/2 pathway as a target of CSF-1R blockade in normoxia (Figure 1E) suggested the possibility of a similar pattern in hypoxia. Indeed, we confirmed this pattern by a gene set enrichment analysis (GSEA) (Figure 3G) and by immunoblotting of phosphorylated sites p-ERK1/2 sites in hypoxia (Figure 3H). Inhibition of ERK1/2 with the MEK1 inhibitor U0126 was able to downregulate the expression of HIF-2α similar to that observed with edicotinib (Figure 3I). In addition, HIF-2α protein expression is a direct sensor of intracellular oxygen availability. Oxygen diffusion from the cellular environment into the cell is primarily controlled by the plasma membrane permeability which is profoundly affected by the membrane concentration of cholesterol (Dotson *et al*, 2017). A decrease in membrane cholesterol will favor oxygen diffusion into the cell. Therefore, the decrease in cholesterol synthesis induced by CSF1R blockade suggested a complementary involvement of the cellular cholesterol in HIF-2α expression. Indeed, the RNAi targeting of SREBF2, which led to a decrease in cholesterol synthesis and cellular uptake, showed a small decrease in HIF-2α (Figure 3J), suggesting a possible combined effect between SREBF2- and ERK1/2-dependent processes. Examination of the expression of cholesterol synthesis genes (*HMGCR*, *DHCR7*) and cellular uptake (*LDLR*) confirms that these two mechanisms are independent as ERK1/2 inhibition did not downregulate these genes, in contrast to edicotinib exposure (Supplementary Figure 3C).

**Figure 3.**
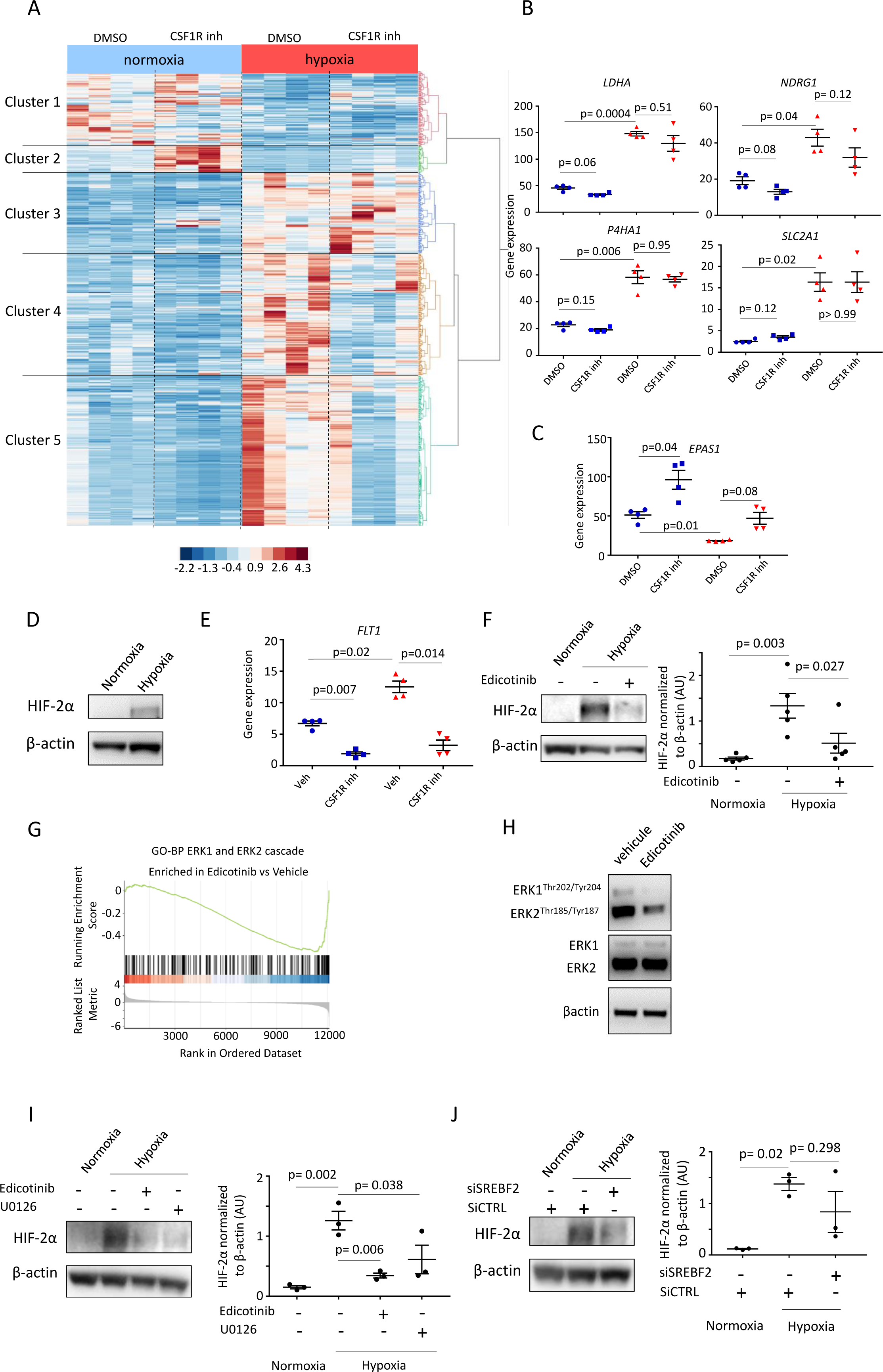
CSF-1R targeting downregulates HIF2α expression in hypoxia through inhibition of ERK1/2 phosphorylation. (A) Hierarchical clustering (Ward methodology) of hypoxic responsive genes (log_2_ FC <-1 and log_2_ FC >-1 and q-value < 0.05) for normoxic and hypoxic macrophages exposed to edicotinib and vehicle (DMSO). (B) Gene expression of *LDHA, NDRG1, P4HA1* and *SLC2A1* genes in normoxic and hypoxic human macrophages treated by edicotinib or vehicle (n=4). Gene expression levels are expressed in FPKM=Fragments Per Kilobase Million. Normoxia is in blue and hypoxia is in red. (C) Gene expression of *EPAS1* gene in normoxic and hypoxic human macrophages treated by edicotinib or vehicle (n=4). Gene expression levels are expressed in FPKM=Fragments Per Kilobase Million. Normoxia is in blue and hypoxia is in red. (D) Immunoblot of HIF-2α in normoxic and hypoxic macrophages (Representative of five independent experiments). (E) Gene expression of *FLT1* gene in normoxic and hypoxic human macrophages treated by edicotinib or vehicle (n=4). Gene expression levels are expressed in FPKM=Fragments Per Kilobase Million. Normoxia is in blue and hypoxia is in red. (F) Immunoblot of HIF-2α in normoxic and hypoxic macrophages treated by vehicle or edicotinib with a representative experiment (left panel) and quantification of the expression of HIF-2α compared to βactin (right panel) (n=4). (G) Gene Set Enrichment Analysis of the Gene Ontology – Biological Process ERK1 and ERK2 cascade for hypoxic macrophages treated by edicotinib. (H) Immunoblot of phosphorylated ERK1/2, total ERK1/2 and βactin in edicotinib treated macrophages (Representative of three independent experiments). (I) Immunoblot of HIF-2α in normoxic and hypoxic macrophages treated by vehicle, edicotinib or U0126 (10µM) with a representative experiment (left panel) and quantification of the expression of HIF-2α compared to βactin (right panel) (n=3). (J) Immunoblot of HIF-2α in normoxic and hypoxic macrophages treated by siRNA against SREBF2 or using siRNA scrambled with a representative experiment (left panel) and quantification of the expression of HIF-2α compared to βactin (right panel) (n=3).

### Hypoxia-induced increase in fatty acid synthesis reversed by CSF-1R targeting in human macrophages

The recent recognition that hypoxia activates SREBP2 in hypoxic THP1-monocytes raised the possibility that the ability of CSF-1R receptor blockade to dampen cholesterol synthesis could be antagonized in hypoxic macrophages (Nakahara *et al*, 2023). However, these authors reported a small or absent effect in hypoxic differentiated THP1-macrophages (Nakahara *et al*, 2023). We sought to address this issue in our primary human macrophages. Examination of differentially repressed genes in exposed hypoxic macrophages confirmed the results obtained in normoxia with a downregulation of the cholesterol synthesis and the ERK1/2 pathway under CSF-1R blockade (Figure 4A). In addition to cholesterol synthesis and cellular uptake under the control of *SREBF2*, we observed that *SREBF1* and *FASN* were similarly downregulated in hypoxia under CSF-1R blockade (Figure 4B). SREBP1, the transcription factor encoded by the *SREBF1* gene, is involved in fatty acid synthesis and fatty acid synthase (FASN) is under the control of SREBPs transcription factors. Furthermore, the analysis of the hypoxia-specific response genes modulated by edicotinib (cluster 4 in Figure 3A) revealed that lipid transport, localization is associated with the hypoxia response (Figure 4C), and that this response could be disrupted by CSF-1R blockade. *FASN*, a gene under the control of *SREBF1* and *SREBF2*, is the major enzymatic complex responsible for the synthesis of palmitic acid (C16:0) and stearic acid (C18:0) from malonyl-CoA, which are subsequently converted to palmitoleic acid (C16:1 cis) and oleic acid (C18:1 cis) respectively (Figure 4D). Fatty acids, particularly oleic acid, have been implicated in macrophage commitment to an immunosuppressive CD206^+^CD38^+^ phenotype in tumor-associated macrophages (Wu *et al*, 2019). Using gas chromatography-mass spectrometry (GC-MS), we confirmed the significant increase of these fatty acids in hypoxic macrophages and the ability of the CSF-1R blockade to attenuate this synthesis (Figure 4E). We also confirmed the downregulation of CD206 expression in tumor-educated macrophages (Figure 4F & Supplementary Figure 4A) and *CD206*, *CD38* gene expression in hypoxic macrophages exposed to edicotinib (Supplementary Figure 4 B). We also found that inhibition of FASN by a specific inhibitor (C75) in hypoxic macrophages was sufficient to prevent IL-4 driven CD206 expression (Figure 4G). These results support for a fatty acid synthase-mediated inhibition of CD206 following CSF-1R targeting.

**Figure 4.**
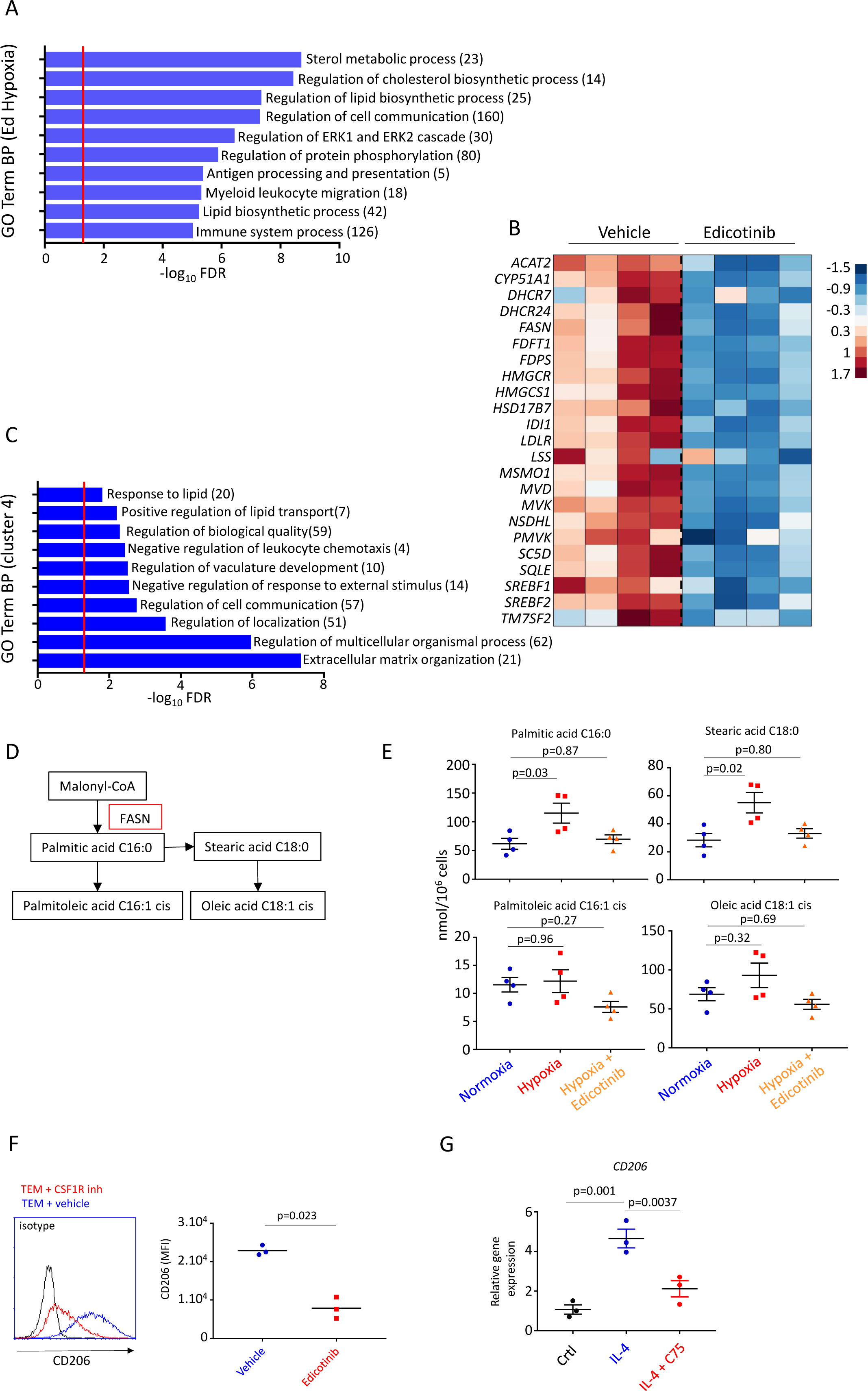
Fatty acid synthesis in hypoxia is reversed by targeting CSF-1R. (A) GO (Gene Ontology)-term analysis of biological processes associated with downregulated genes under edicotinib exposure in hypoxia. Number of genes associated with each GO-terms are indicated. (B) Heat map of cholesterol synthesis genes associated with *SREBF1*, *FASN* and *SREBF2* for hypoxic macrophages treated by edicotinib compared to vehicle. (C) GO (Gene Ontology)-term analysis of biological processes associated with genes identified in cluster 4 of figure 3A. Number of genes associated with each GO-terms are indicated. (D) Fatty acid synthesis pathway involving the fatty acid synthetase (FASN). (E) Quantification by GS-MS of palmitic acid (C16:0), stearic acid (C18:0), palmitoleic acid (C16:1 cis) and oleic acid (C18:1 cis) in macrophages in normoxia, in hypoxia and in hypoxia treated by edicotinib (n=4). (F) Quantification by flow cytometry of CD206 membrane expression in co-cultured macrophages with RKO cells and IL-4 (25 ng/mL) for 48 h with vehicle (DMSO) or edicotinib (n=3). Representative panel on the left and quantification on the right. (G) Expression of *CD206 (MRC1)* gene in hypoxic macrophages stimulated by IL-4 for 4 hours and a specific inhibitor of FASN (C75 50 µM) determined by RT-qPCR (n=3).

### CSF-1R targeting downregulates the expression of dihydropyrimidine dehydrogenase preventing 5-FU hypoxic macrophage-driven chemoresistance

HIF-2α has been recognized as an important regulator of innate immunity in the context of tumors, in particular by controlling macrophage’s recruitment (Imtiyaz *et al*, 2010). In addition, we recently reported that HIF-2α controls the expression of dihydropyrimidine dehydrogenase (DPD) in hypoxia independently of its transcription factor activity. DPD expression in hypoxic macrophages is responsible for a resistance to 5-fluorouracil in human colorectal cancer (Malier *et al*, 2021). We have shown in the same study that rodent models are inadequate to mimic this mechanism, because the DPYD gene is epigenetically negatively regulated in rodent macrophages, in contrast to humans. Downregulation of HIF-2α expression under CSF-1R blockade offers a potential strategy to restore chemosensitivity in this context. Indeed, edicotinib is able to downregulate the expression of DPD in hypoxic macrophages (Figure 5A), resulting in the complete abrogation of its enzymatic activity, preventing the conversion of uracil to dihydrouracil (Figure 5B). We confirmed the inhibition of the HIF-2α-dependent synthesis of DPD during the transition to hypoxia, which requires the continuous destabilization of HIF-2α during the transition (Figure 5C). The previous demonstration that ERK1/2 phosphorylation controls HIF-2α expression implies an expected control of DPD expression in the same context as we confirmed (Figure 5D) as well as the implication of the cellular cholesterol content in the hypoxia-induced DPD expression in macrophages (Figure 5E). Consequently, the downregulation of DPD expression obtained by CSF1R targeting allows the restoration of 5-FU sensitivity of HT-29 and RKO colorectal cancer cells in a co-culture setting (Figure 5F). We have previously shown that DPD expression under the HIF-2α expression is not related to mRNA modulation (Malier *et al*, 2021), to ensure that this is still the case in the tumor environment we showed that edicotinib downregulates DPD expression in hypoxic TAMs co-cultured with tumors (Figure 5G) without downregulating its mRNA as expected (Figure 5H). We have previously shown that macrophages are the major source of DPD expression in colorectal cancer (Malier *et al*, 2021). However, these macrophages are replenished by circulating monocytes and these hypoxic monocytes (CD14+) express DPD in contrast to other immune cells such as hypoxic T (CD3+) or B (CD19+) lymphocytes (Figure 5I & Supplementary Figure 5), representing another source of immune-driven chemoresistance to 5-FU. Circulating monocytes are recruited to tumor sites by CCL2/CCL7 chemokine secretion from tumor macrophages (Qian *et al*, 2011; Sanford *et al*, 2013; Yang *et al*, 2020). Edicotinib may interfere with this process. Indeed, the gene ontology – biological process associated with myeloid leukocyte migration was associated with differentially downregulated genes by CSF-1R blockade (Figure 4A). We confirmed this pattern by the effect of CSF-1R targeting on the expression of *CCL2* and *CCL7* in macrophages (Figure 5J). Similarly, CCL2 and CCL7 expressions are disrupted in hypoxic TAMs exposed to edicotinib (Figure 5K). These results demonstrate the ability of CSF-1R targeting to disrupt DPD expression in tumor-associated macrophages and prevent the recruitment of DPD-expressing myeloid cells to the tumor site.

**Figure 5.**
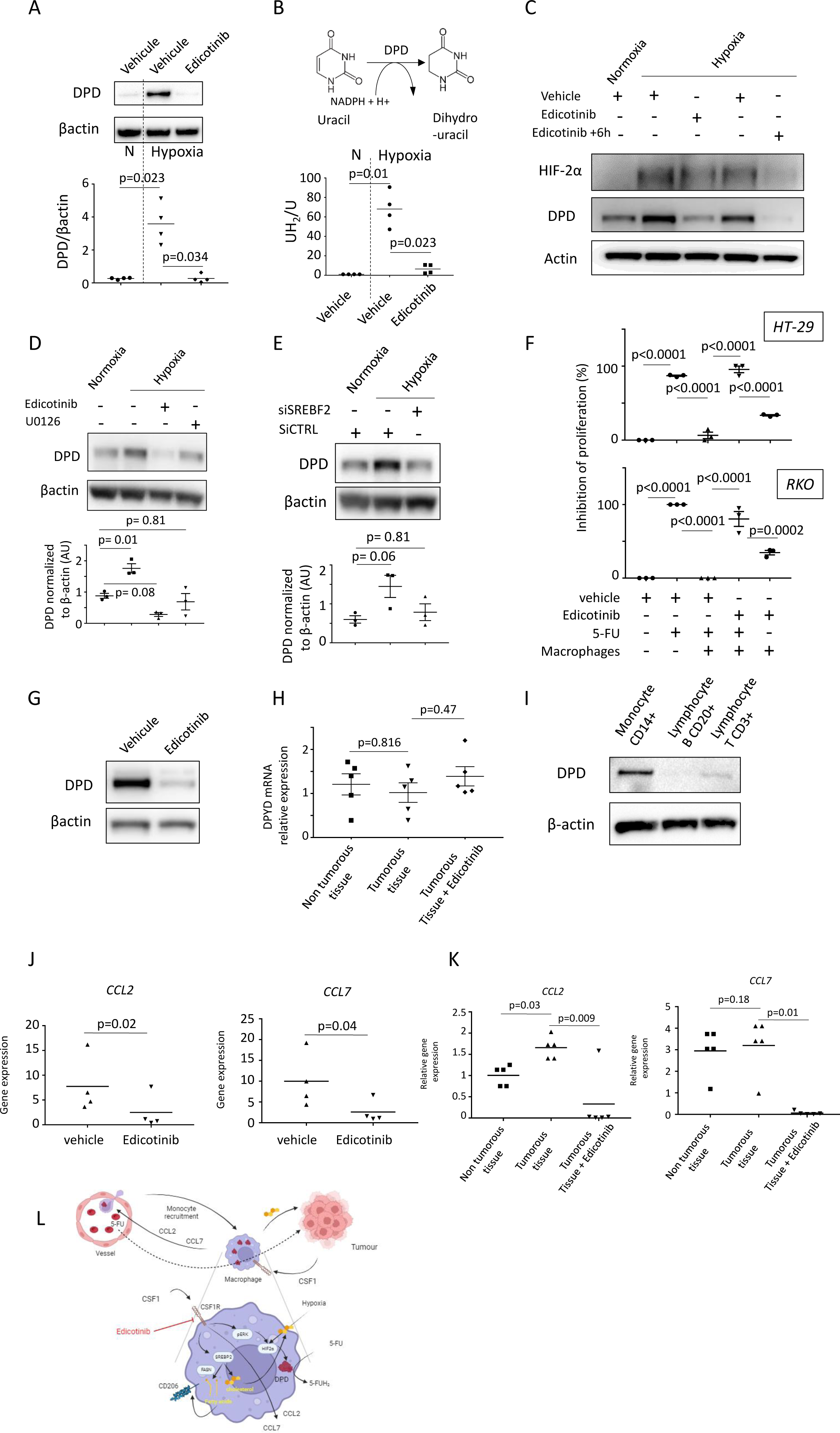
CSF-1R targeting downregulates dihydropyrimidine dehydrogenase expression in tumor-associated macrophages. (A) Immunoblot of DPD in normoxic and hypoxic macrophages treated by vehicle (DMSO) and edicotinib with a representative experiment (upper panel) and quantification of the expression of HIF-2α compared to βactin (lower panel) (n=4). (B) Quantification of the reductive activity of DPD through the conversion of uracil (U) to dihydrouracil (UH_2_) (n=4). (C) Immunoblot of HIF-2α, DPD and βactin for macrophages treated by edicotinib for 48h then submitted to 6h hypoxia with or without maintaining the CSF-1R blockade (+6h condition). Representative of three independent experiments. (D) Immunoblot of DPD and βactin for macrophages treated by edicotinib or U0126 for 48h then submitted to 6h hypoxia with edicotinib or U0126. A representative experiment is shown in the upper panel and the quantification compared to βactin appears in the lower panel (n=3). (E) Immunoblot of DPD and βactin for macrophages treated by siRNA against SREBF2 or SiRNA scrambled for 48h then submitted to 6h hypoxia. A representative experiment is shown in the upper panel and the quantification compared to βactin appears in the lower panel (n=3). (F) Co-culture between colorectal cancers cells (HT-29, RKO) with human THP1-macrophages. 5-FU was used at 1µg/mL for 48h. Edicotinib (5µM) or vehicle (DMSO) was added during 48h (n=3). (G) Immunoblot of DPD and β-actin in hypoxic tumor associated macrophages treated by edicotinib or vehicle (DMSO) (representative of three experiments). (H) Expression of *DPYD* gene in hypoxic macrophages cultured with non-tumorous and tumorous tissues determined by RT-qPCR. Tumor associated macrophages were also treated by edicotinib (n=5). Relative expression compared to control macrophages in normoxia. (I) Immunoblot of DPD and β-actin in CD14+ monocytes, CD3+ T lymphocytes and CD20+ B lymphocytes (representative of three independent healthy donors). (J) Gene expression of *CCL2* and *CCL7* in hypoxic human macrophages treated by edicotinib (n=4). Gene expression levels are expressed in FPKM=Fragments Per Kilobase Million. (K) Expression of *CCL2* and *CCL7* genes in hypoxic macrophages cultured with non-tumorous and tumorous tissues determined by RT-qPCR (n=5). Tumor associated macrophages were also treated by edicotinib. Relative expression compared to control macrophages in normoxia. (L) Schematics of CSF-1 controlled tumor associated macrophages involvement in tumor growth and consequences of CSF-1R targeting. Figure created using Biorender (biorender.com)

## Discussion

Tumor-associated macrophages are an essential component of the tumor immune microenvironment in the vast majority of cancers. Their involvement in angiogenesis, extracellular matrix remodeling, cancer cell proliferation, metastasis spreading and treatment resistance is leading to the development of macrophage-centered therapeutic strategies (Mantovani *et al*, 2022). The importance of the CSF-1/CSF-1R axis in macrophage survival, differentiation and activation makes CSF-1R targeting a favored approach to remodel TAMs. Two strategies have been pursued to achieve this goal: blocking the interaction between the ligand (CSF-1, IL-34) and its receptor using specific anti-CSF-1R antibodies or inhibiting the tyrosine kinase activity of CSF-1R using specific inhibitors. Various inhibitors have been developed over the years, and despite high specificity against CSF-1R, these molecules have failed to show a strong clinical effect when used as monotherapy (El-Gamal *et al*, 2018; Mantovani *et al*, 2022). The unclear reasons for this lack of efficacy led us to design the current study to investigate in detail the molecular consequences of CSF-1R blockade using a selective CSF-1R kinase inhibitor (edicotinib or JNJ40346527), which has already been validated in humans (Genovese *et al*, 2015; von Tresckow *et al*, 2015).

We observe a clear metabolic shift in CSF-1R-targeted macrophages, mainly focused on cholesterol and fatty acid metabolism. The link between CSF-1 and cholesterol metabolism was suggested in a previous study (Irvine *et al*, 2009). We confirm the ability of CSF-1 to induce cholesterol synthesis and show that CSF-1R blockade is able to downregulate *SREBF2* expression leading to the inhibition of the cholesterol synthesis but also its cellular uptake. We observe that colorectal tumors favor *SREBF2* expression and consequently drives the increase of cholesterol levels in human macrophages. Cholesterol efflux from macrophages under the control of transporter proteins such as ABCA1/ABCG1 has been associated with tumor growth (Goossens *et al*, 2019). Tumor cells exposed to macrophage-secreted cholesterol secrete hyaluronic acid that drives macrophage reprogramming toward an M2 phenotype in mice (Goossens *et al*, 2019). Even if cholesterol transporters are not suppressed in macrophages targeted by edicotinib, the decreased total amount of cellular cholesterol induced by edicotinib prevents the ability of macrophages to maintain their secretion, thereby braking the feedback loop between tumor-associated macrophages and tumors.

Since cholesterol is synthesized by the metabolic synthesis starting from acetyl-coA, which is produced by the fatty acid β-oxidation pathway, we investigated the fatty acid synthesis in macrophages. We find that edicotinib is able to prevent the increase in fatty acid accumulation driven by hypoxia in human macrophages (Figure 4E), thereby preventing the appearance of an immunosuppressive phenotype CD206^+^CD38^+^ induced by oleate (Supplementary Figure 4).

The CSF-1R blockade strategy is designed to occur in the low-oxygen tumor environment. The sensitivity of cholesterol synthesis and CSF-1 coupling in hypoxia is not known. A recent study highlighted the importance of hypoxia in the induction of cholesterol synthesis in monocytes but not in differentiated macrophages (Nakahara *et al*, 2023). The influence of CSF-1R targeting on the hypoxic response shows that *EPAS1,* which encodes HIF-2α, is upregulated. In conjunction with this observation, we note the inverse correlation between *EPAS1* mRNA and HIF-2α protein, revealing the counterregulation of *EPAS1* transcription by HIF-2α itself. A similar observation has been proposed for HIF-1α, based on miR-429, whose expression is promoted by HIF-1α (Bartoszewska *et al*, 2015). However, this negative feedback loop was associated with the decrease of HIF-1α in chronic hypoxia. Our observation was made in chronic hypoxia and no decrease of HIF-2α protein was observed. This observation needs to be further investigated to determine the direct or indirect mechanism. Our observation of the association between CSF-1R targeting and HIF-2α expression revealed the role of ERK1/2 activation in stabilizing this hypoxia-inducible factor in human macrophages. The downregulation of HIF-2α opens a promising avenue to target the recently identified mechanism leading to the chemoresistance to 5-FU driven by hypoxic macrophages in colorectal cancer (Malier *et al*, 2021). In this study, we show that targeting CSF-1R downregulates ERK1/2 phosphorylation, which in turn prevents HIF-2α expression in hypoxia, thereby preventing the expression of DPD, which can no longer degrade 5-fluorouracil. We further find that monocyte recruitment driven by the CCL2/CCL7 axis is also disrupted, preventing the contribution of other myeloid DPD-expressing cells to tumor infiltration.

In this study, we have provided compelling evidence showing that CSF-1R blockade is an effective strategy to prevent cancer cell-driven TAM recruitment and macrophage-induced resistance to chemotherapy. We have demonstrated the efficacy of an inhibitor of CSF-1R kinase activity to achieve metabolic reprogramming of tumor-associated macrophages by interfering with the cholesterol and fatty acid mediated cross-talk between macrophages and cancer cells.

## Material and Methods

### Resources

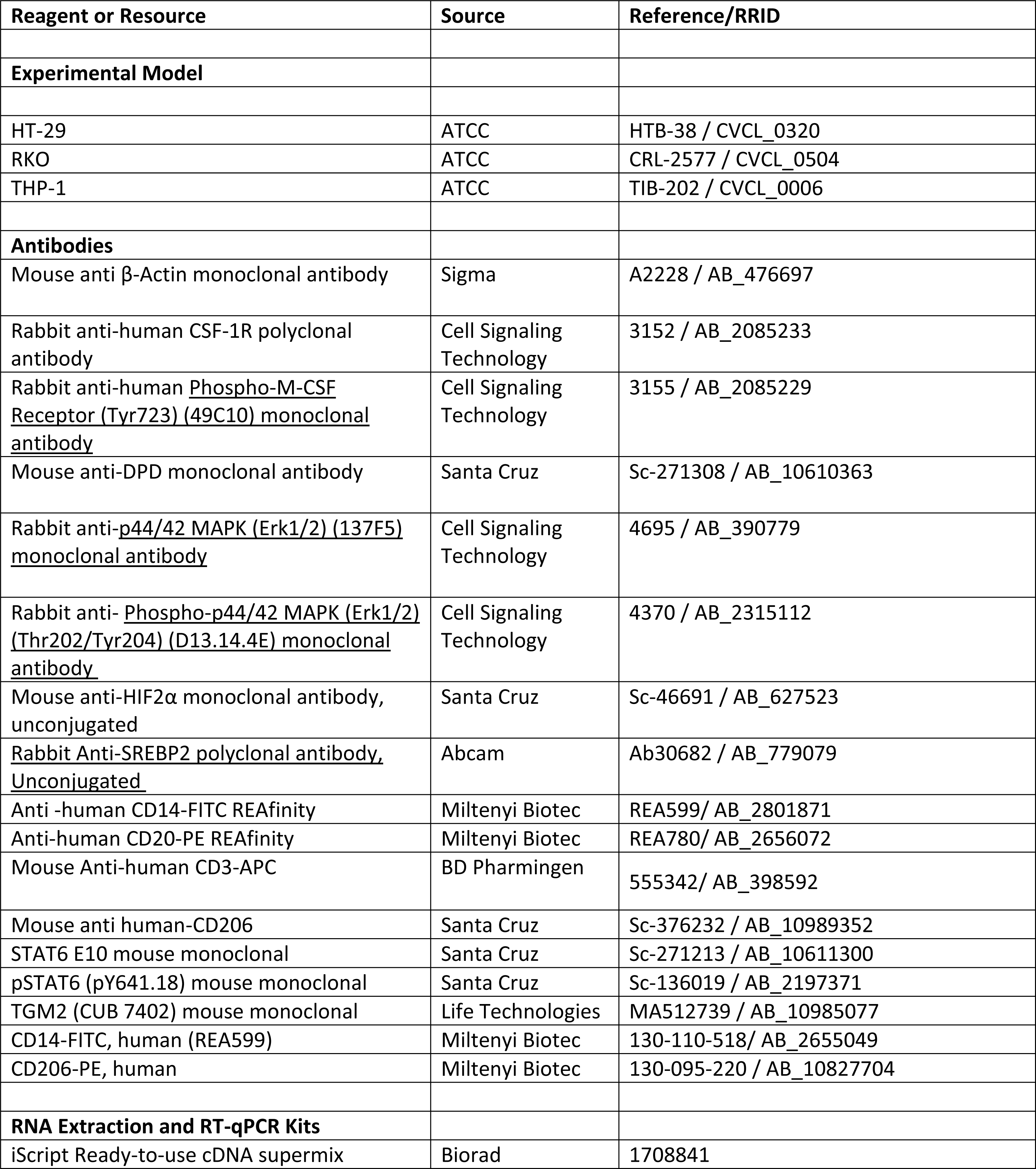
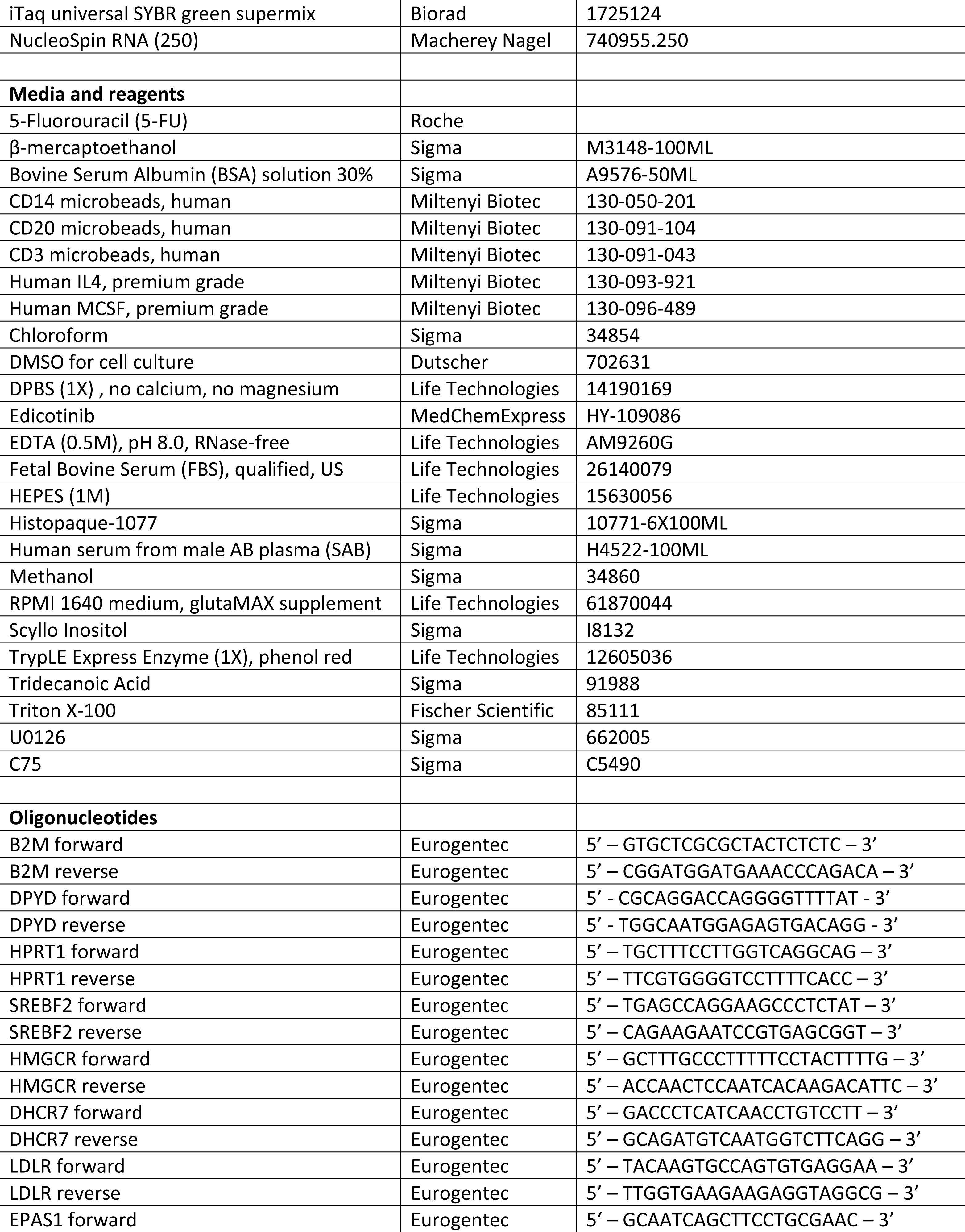
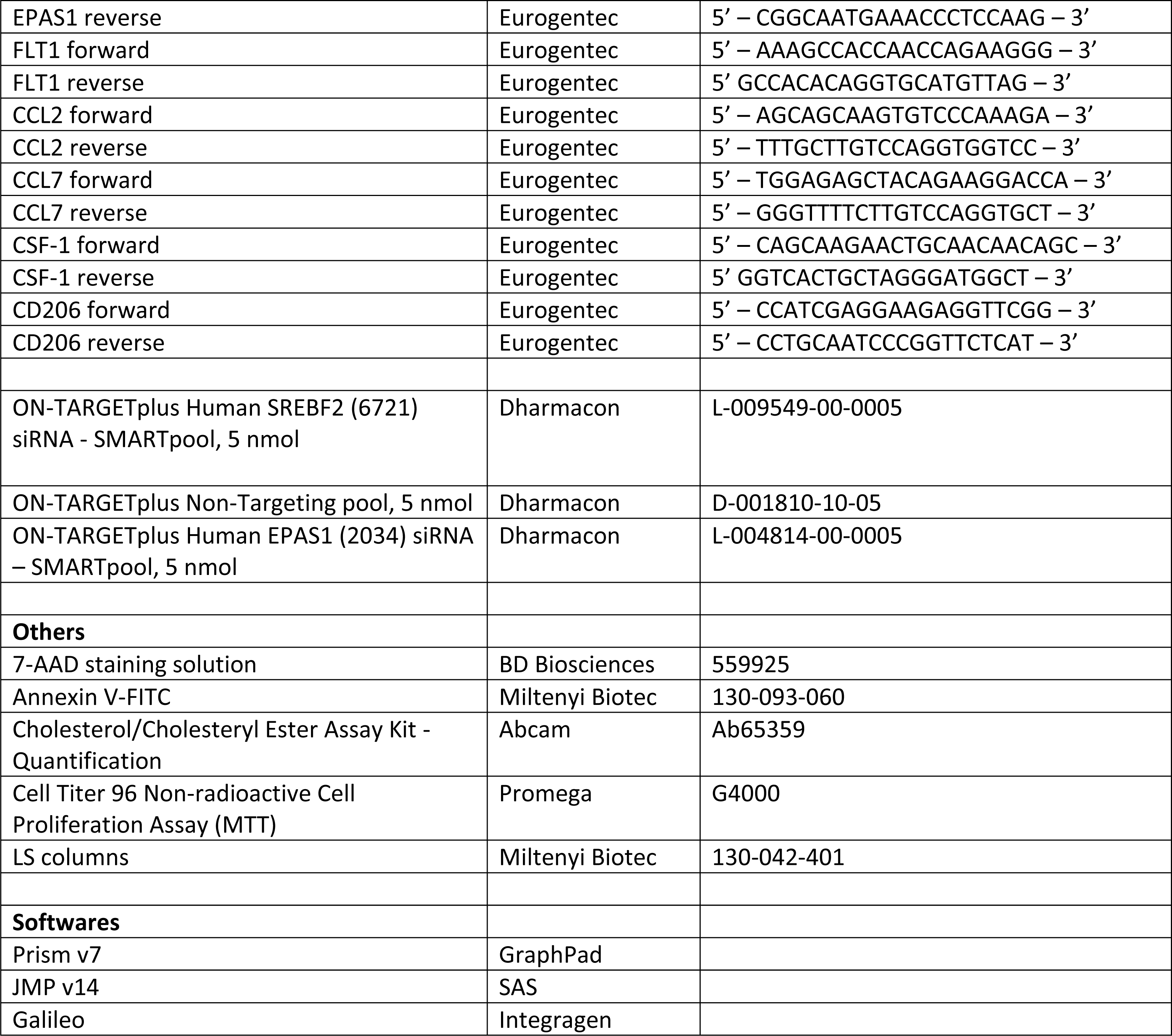

### Human samples

Human blood samples from healthy de-identified donors were obtained from EFS (French national blood service) as part of an authorized protocol (CODECOH DC-2018–3114). Donors gave signed consent for use of their blood in this study. Tumorous and non tumorous tissues were obtained from surgically resected colons at the university hospital of Grenoble Alpes from patients included in the CRC-ORGA2 clinical trial (clinicaltrial.gov, NCT05038358). Inclusion criteria comprise adult patients suffering from colorectal adenocarcinoma treated by surgery without neoadjuvant chemotherapy (Table).

| Sexe | Age | Histologic diagnosis | Surgery | TNM | Molecular charactrization |
| --- | --- | --- | --- | --- | --- |
| F | 92 | Adenocarcinoma | Right colectomy (coelioscopic) | pT4N1R0 | p-MMR |
| M | 71 | Adenocarcinoma poorly differentiated | Right colectomy (coelioscopic) | pT3N1R0 | MSI (loss MSH6) |
| F | 55 | Adenocarcinoma poorly differentiated low grade | Sigmoidectomy | pT3N2bR0 | NA |
| F | 69 | adenocarcinoma well differnetiated on tubulovillous adenoma | Left Colectomy | pT3pN1R0 | pMMR |
| M | 74 | Adenocarcinoma (lieberkühnien) | Left Colectomy | pT3N0 | MSS |
| F | 70 | Adenocarcinoma (lieberkühnien) low grade | Left colectomy | pT3N0M0 | MMR |
| M | 63 | Adénocarcinoma low grade | Right colectomy (coelioscopic) | pT4N1b | MMR |
| M | 84 | Adenocarcinoma (lieberkühnien) low grade | Right colectomy (coelioscopic) | pT3N0R0 | NA |

### Cell culture

THP-1, RKO and HT-29, were purchased from ATCC. THP-1, HT-29 and RKO were maintained in RPMI (Gibco) supplemented with 10% FBS (Gibco) at 37°C. All cells were routinely tested for mycoplasma contamination using MycoAlert detection kit (Lonza).

### Human macrophage differentiation from monocytes

Monocytes were isolated from leukoreduction system chambers of healthy EFS donors using differential centrifugation (Histopaque 1077, Sigma) to obtain PBMCs. CD14^+^ microbeads (Miltenyi Biotec) were used to select monocytes according to the manufacturer’s instructions. Monocytes were plated in RPMI (Life Technologies) supplemented with 10% SAB (Sigma), 10 mM HEPES (Life Technologies), MEM Non-essential amino acids (Life Technologies) and 25 ng/mL M-CSF (Miltenyi Biotec). Differentiation was obtained after 6 days of culture. Hypoxic cultures were performed in a hypoxic chamber authorizing an oxygen partial pressure control (HypoxyLab, Oxford Optronix, UK). Hypoxia experiments were performed at 25 mmHg of oxygen (∼ 3%). CD3^+^ T lymphocytes and CD20^+^ B lymphocytes were sorted from PBMCs using CD3^+^ and CD20^+^ microbeads (Miltenyi Biotec) according to the manufacturer’s instructions.

### Preparation of tumor associated macrophages

Tumorous and non-tumorous tissues obtained from colon of CRC patients. Tissues were kept and transported in ice in RPMI medium supplemented with non-essential amino acids and HEPES as well as antibiotics including penicillin-streptomycin and gentamicin and antifungal amphotericin B. Tissues were minced in pieces of 2-4 mm^3^ and deposited on cell culture inserts (4 pieces by insert). These inserts were deposited in 6 well plates on 2 mL of RPMI + 10% human serum for 24 h. The resulting medium produced by tumorous and non-tumorous tissues was then used to culture human monocyte derived macrophages for 48h to obtain tumor-associated macrophages.

### Macrophage Conditioned Medium (MCM)

THP1 macrophages were plated (12-wells plates) and differentiated at 500,000 cells with 50nM of PMA for 24 hours. THP1 macrophages were cultured at 3% oxygen and treated with Edicotinib at 3µM or DMSO acting as a vehicle for 48 hours. Then 1µg/mL of 5-FU was used to generate the 5FU Edicotinib MCM or 5FU Vehicle MCM during 24h. MCM was added for 48 hours to two distinct colon cancer cell lines, RKO and HT-29 cells, which were plated previously at 100,000 cells per well (12-wells plates) for 24 hours. Then cancerous cells were collected and counted and the percentage of growth inhibition was assessed for each condition.

### RNAseq

RNA extraction was performed using the NucleoSpin RNA kit components (Macherey Nagel) according to the manufacturer’s instructions. RNA sequencing was performed using an Illumina HiSeq 4000 sequencer (Integragen). Gene expression quantification was performed using the STAR software. STAR obtains the number of reads associated to each gene in the Gencode v31 annotation (restricted to protein-coding genes, antisense and lincRNAs). Raw counts for each sample were imported into R statistical software. Extracted count matrix was normalized for library size and coding length of genes to compute FPKM expression levels. The Bioconductor edgeR package was used to import raw counts into R statistical software. Differential expression analysis was performed using the Bioconductor limma package and the voom transformation. Gene ontology analysis was performed using the GONet software (https://tools.dice-database.org/GOnet/). Gene list from the differential analysis was ordered by decreasing log2 fold change. Gene set enrichment analysis (GSEA) was performed by clusterProfiler::GSEA function using the fgsea algorithm.

### Lipidomics

Human macrophages were metabolically quenched in dry ice – ethanol for one minute then washed three times in cold PBS. Then total lipids were extracted in chloroform/methanol/water (1:3:1, v/v/v) containing PC (C13:0/C13:0), 10 nmol and C21:0 (10 nmol) as internal standards for extraction) for 1 h at 4◦C, with periodic sonication. Then polar and apolar metabolites were separated by phase partitioning by adding chloroform and water to give the ratio of Chloroform/Methanol/Water, 2:1:0.8 (v/v/v). For lipid analysis, the organic phase was dried under N2 gas and dissolved in 1-butanol to obtain1µl butanol/10^7^ macrophages.

*Total lipid analysis* – Total lipid was then added with 1 nmol pentadecanoic acid (C15:0) as internal standard and derivatized to give fatty acid methyl ester (FAME) using trimethylsulfonium hydroxide (TMSH, Machenery Nagel) for total glycerolipid content. Resultant FAMEs were then analysed by GC-MS as previously described (Dubois *et al*, 2018). All FAME were identified by comparison of retention time and mass spectra from GC-MS with authentic chemical standards. The concentration of FAMEs was quantified after initial normalization to different internal standards and finally to macrophage number.

### Immunoblotting

Cells were lyzed in RIPA buffer supplemented with protease inhibitors (AEBSF 4 mM, Pepstatine A 1 µM and Leupeptin 0.4 mM; Sigma Aldrich) and HIF-hydroxylase inhibitor (DMOG 1 mM, Sigma Aldrich) for total lysate or direct lysis in Laemmli buffer (2X). Proteins were quantified by BCA assay (ThermoFischer) and 15 µg of total protein were run on SDS-PAGE gels. Proteins were transferred from SDS-PAGE gels to PVDF membrane (Biorad), blocked with TBS-Tween supplemented with 5% milk, primary antibodies were incubated at 1 µg/mL overnight 4°C. After washing with TBS, the membrane was incubated with a horseradish peroxidase-conjugated secondary antibody (Jackson Immunoresearch). Signal was detected by chemoluminescence (Fusion FX imaging system, Vilber) after exposition to Clarity ECL substrate (Biorad).

### Flow cytometry

Flow cytometry data was acquired on an Accuri C6 (BD) flow cytometer. Doublet cells were gated out by comparing forward scatter signal height (FSC-H) and area (FSC-A). At least 10,000 events were collected in the analysis gate. Median fluorescence intensity (MFI) was determined using Accuri C6 software (BD).

### DPD activity measurements

These analyses were performed in the Pharmacology Laboratory of Institut Claudius-Regaud (France) using an HPLC system composed of Alliance 2695 and diode array detector 2996 (Waters). Uracil (U), Dihydrouracil (UH2), Ammonium sulfate 99%, Acetonitrile (ACN) gradient chromasolv for HPLC and 2-propanol were purchased from Sigma. Ethyl acetate Scharlau was of HPLC grade and purchased from ICS (Lapeyrouse-Fossat, France). Water from Milli-Q Advantage A10 and MultiScreen-HV 96-well Plates were used (Merck Millipore). Calibration ranges were 3.125 – 200 ng/mL for U and 25 - 500 ng/mL for UH2 and 5-FU (5µg/mL) was used as an internal standard.

### RNA isolation and qPCR analysis for gene expression

Cells were directly lyzed and RNA was extracted using the NucleoSpin RNA kit components (Macherey Nagel) according to the manufacturer’s instructions. Reverse transcription was performed using the iScript Ready-to-use cDNA supermix components (Biorad). qPCR was then performed with the iTaq universal SYBR green supermix components (Biorad) on a CFX96 (Biorad). Quantification was performed using the ΔΔCt method using two different housekeeping genes.

### Quantification of Cholesterol

Macrophages were treated with 10 µM of Edicotinib or Simvastatin for 72 hours afterwhich cells and supernatants were retrieved. Supernatants were stored at -80°C. Lipids were extracted from cells using choloroform:isopropanol:NP-40 (7:11:0.1). Organic phase was taken and air-dried at 50°C and stored at -80°C. The amount of cholesterol in cells and supernatants was assessed using the cholesterol/cholesteryl ester assay quantification kit (ab65359, abcam) according to the manufacturer’s instructions.

### Silencing of EPAS1 and SREBF2

Fully differentiated macrophages were transfected with siEPAS1 (L-004814-00-0005, Dharmacon) or siSREBF2 (L-009549-00-0005, Dharmacon) at a final concentration of 5nmol/L using Lipofectamine (RNAiMAx, Life Technologies).

### Statistical analysis

Statistics were performed using Graph Pad Prism 7 (Graph Pad Software Inc). When two groups were compared, we used a paired or unpaired two-tailed student’s t-test when appropriate. When more than two groups were compared, we used a one-way ANOVA analysis with Sidak’s multiple comparison test on paired data when appropriate. The likelihood of data according to a null-hypothesis (p-value) is presented in the figures. The number of independent experiments used to generate the data is shown on each figure.

### Data Availability

The data generated and analyzed will be made from the corresponding author on reasonable request.

## Acknowledgments

KG has been supported by the ITN (International Training Network) Phys2Biomed project, which was funded from the European Union’s Horizon 2020 research and innovation programme under the Marie Skłodowska-Curie grant agreement No 812772 and is supported by the ERiCAN program of Fondation MSD-Avenir (Reference DS-2018-0015). CGP is supported by the Innovative Research Initiative (IRGA 2022) of the university Grenoble Alpes. MM is supported by the APMC Fondation (Agir pour les Maladies Chroniques). This work is supported by the Ligue Régionale contre le Cancer. CYB, and YYB are supported by Agence Nationale de la Recherche, France (Project ApicoLipiAdapt grant ANR-21-CE44-0010 ; Project Apicolipidtraffic grant ANR-23-CE15-0009-01), The Fondation pour la Recherche Médicale (FRM EQU202103012700), Laboratoire d’Excellence Parafrap, France (grant ANR-11-LABX-0024), LIA-IRP CNRS Program (Apicolipid project), the Université Grenoble Alpes (IDEX ISP Apicolipid) and Région Auvergne Rhone-Alpes for the lipidomics analyses platform (Grant IRICE Project GEMELI), Collaborative Research Program Grant CEFIPRA (Project 6003-1) by the CEFIPRA (MESRI-DBT). We thank Malika Yakoubi for her technical support on DPD chromatographic activity measurment. We thank Dr Lemoigne, Dr Federspiel and Dr Plasse from the University Hospital of Grenoble-Alpes for kindly providing clinical grade 5-Fluorouracil.

## Contributions

KG, CGP, MM, CH, RG, AM performed experiments and analysed the data. AM supervised the data analysis. FT performed chromatography measurements. YB, CB performed the lipidomics analysis. GR, EG recruited patients. MHL performed the pathological analysis. AM wrote the original draft and all authors were involved in manuscript editing. Funding acquisition and supervision of the study was performed by AM.

## Supplementary Figure Legends

**Supplementary Figure 1.**
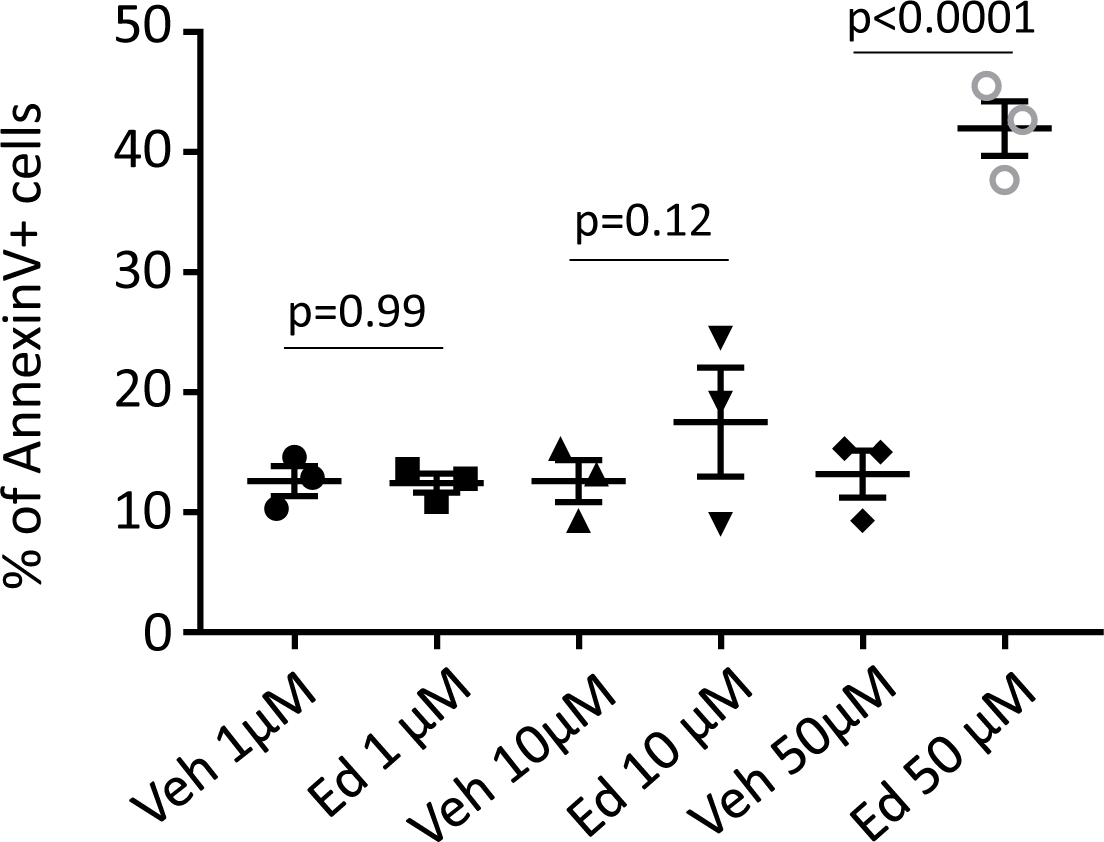
Flow cytometry analysis of Annexin-V+ cells treated for 48h. Each condition has its own vehicle control (DMSO) at the same concentration than edicotinib (Ed) (n=3).

**Supplementary Figure 2.**
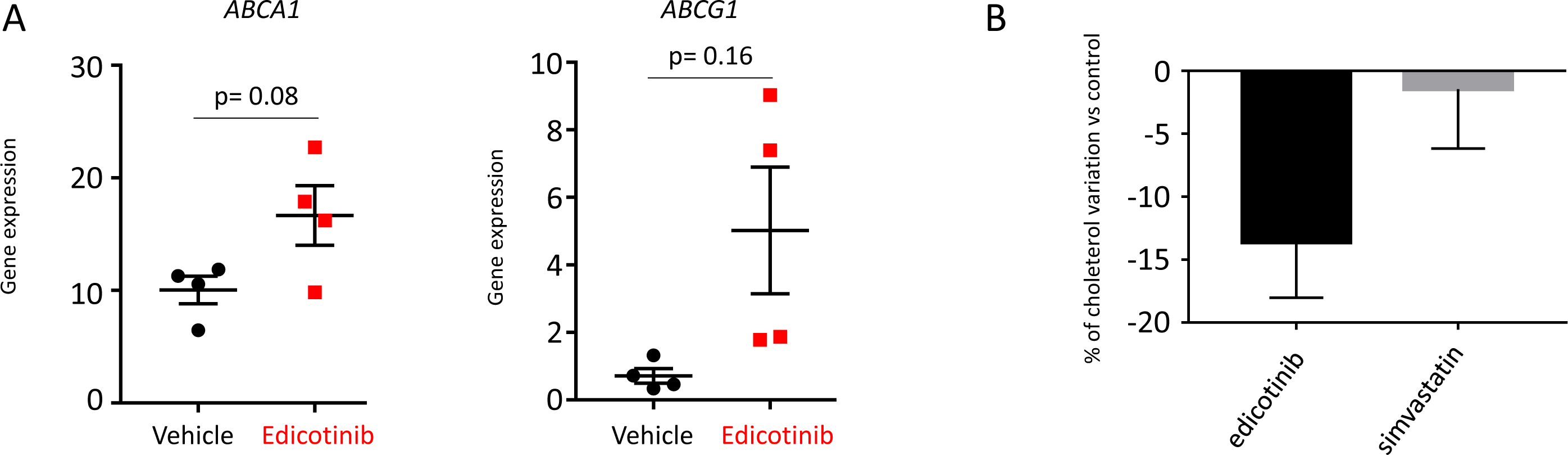
Gene expression of *ABCA1* and *ABCG1* genes in human macrophages treated by edicotinib (n=4). Gene expression levels are expressed in FPKM=Fragments Per Kilobase Million. (A) Quantification of extracellular cholesterol secreted by macrophages treated by edicotinib or simvastatin compared to vehicle (DMSO) (n=3). The amount of decrease is represented in % compared to the control.

**Supplementary Figure 3.**
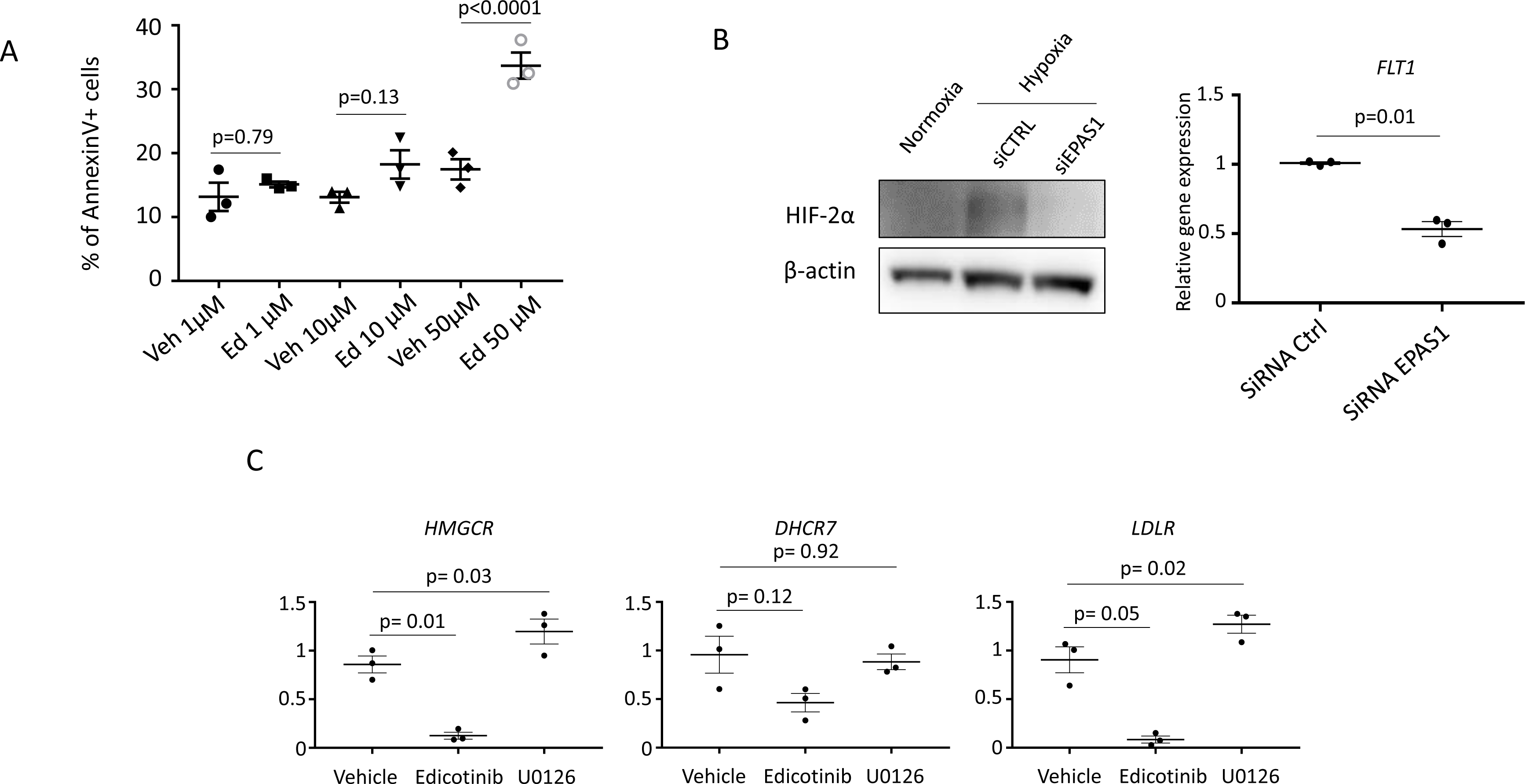
(A) Flow cytometry analysis of Annexin-V+ cells treated for 48h in hypoxia. Each condition has its own vehicle control (DMSO) at the same concentration than edicotinib (Ed) (n=3). (B) Immunoblot of HIF-2α in macrophages in hypoxia (6h) treated by siRNA against SREBF2 or using siRNA scrambled compared to normoxia (representative of three independent experiments, left panel). Expression of *FLT1* gene after 6h in hypoxia in the same macrophages than in the left panel determined by RT-qCPR (right panel) (n=3). (C) Expression of *HMGCR*, *DHCR7* and *LDLR* genes in hypoxic macrophages treated by edicotinib or U0126 (10 µM) or vehicle determined by RT-qPCR (n=3). Relative expression compared to control macrophages in normoxia.

**Supplementary Figure 4.**
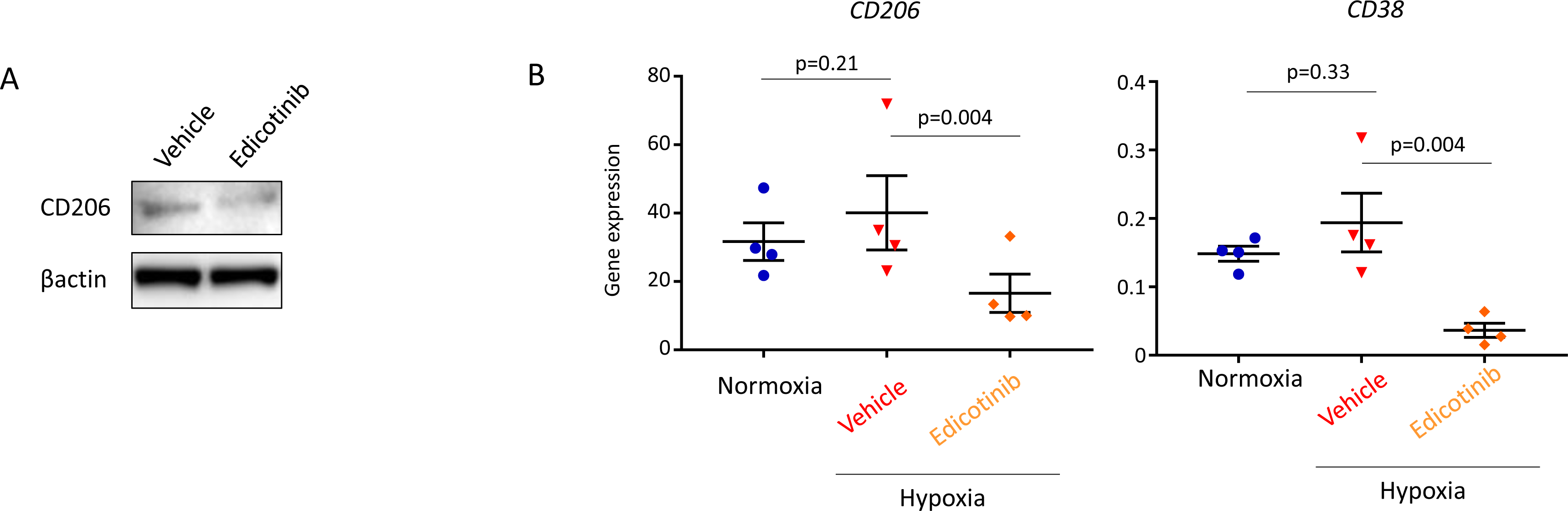
(A) Immunoblot of CD206 in hypoxic macrophages co-cultured with tumors exposed to edicotinib vs vehicle (DMSO) for 24h. Representative of three independent experiments. (B) Gene expression of *CD206* and *CD38* genes in human macrophages treated by edicotinib in hypoxia. Gene expression levels are expressed in FPKM=Fragments Per Kilobase Million.

**Supplementary Figure 5.**
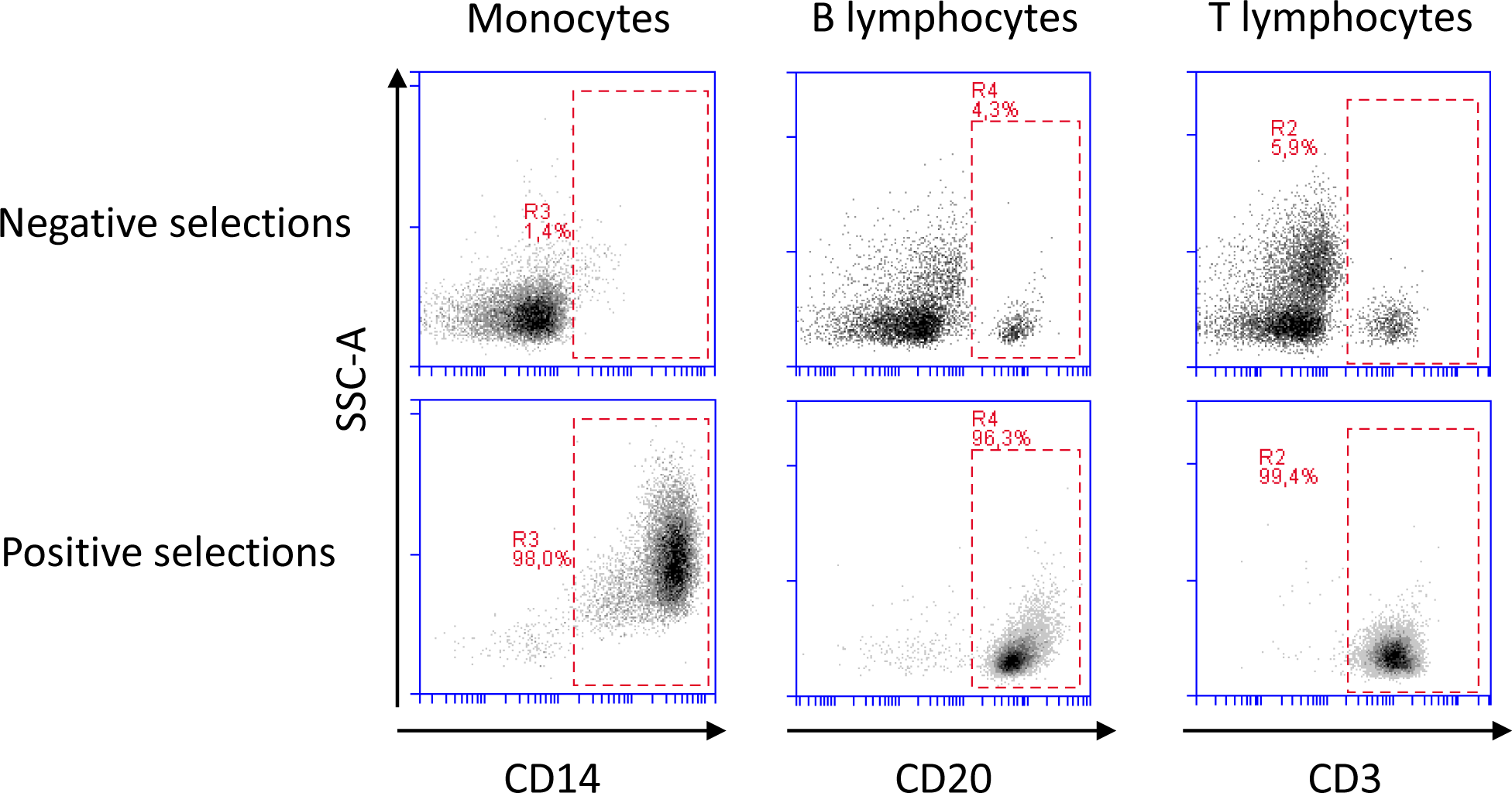
Flow cytometry analysis of sorted Monocytes, T lymphocytes and B lymphocytes using CD14, CD3 and CD20 expression respectively. Representative of three independent experiments on healthy donors.

